# Proteomics Standards Initiative Extended FASTA Format (PEFF)

**DOI:** 10.1101/624494

**Authors:** Pierre-Alain Binz, Jim Shofstahl, Juan Antonio Vizcaíno, Harald Barsnes, Robert J. Chalkley, Gerben Menschaert, Emanuele Alpi, Karl Clauser, Jimmy K. Eng, Lydie Lane, Sean L. Seymour, Luis Francisco Hernández Sánchez, Gerhard Mayer, Martin Eisenacher, Yasset Perez-Riverol, Eugene A. Kapp, Luis Mendoza, Peter R. Baker, Andrew Collins, Tim Van Den Bossche, Eric W. Deutsch

## Abstract

Mass spectrometry-based proteomics enables the high-throughput identification and quantification of proteins, including sequence variants and post-translational modifications (PTMs), in biological samples. However, most workflows require that such variations be included in the search space used to analyze the data, and doing so remains challenging with most analysis tools. In order to facilitate the search for known sequence variants and PTMs, the Proteomics Standards Initiative (PSI) has designed and implemented the PSI Extended FASTA Format (PEFF). PEFF is based on the very popular FASTA format but adds a uniform mechanism for encoding substantially more metadata about the sequence collection as well as individual entries, including support for encoding known sequence variants, PTMs, and proteoforms. The format is very nearly backwards compatible, and as such, existing FASTA parsers will require little or no changes to be able to read PEFF files as FASTA files, although without supporting any of the extra capabilities of PEFF. PEFF is defined by a full specification document, controlled vocabulary terms, a set of example files, software libraries, and a file validator. Popular software and resources are starting to support PEFF, including the sequence search engine Comet and the knowledge bases neXtProt and UniProtKB. Widespread implementation of PEFF is expected to further enable proteogenomics and top-down proteomics applications by providing a standardized mechanism for encoding protein sequences and their known variations. All the related documentation, including the detailed file format specification and example files, are available at http://www.psidev.info/peff.

## Introduction

Mass spectrometry (MS) based proteomics has become the most commonly used technique for detecting the presence of and measuring the abundance of proteins in biological samples^1^. Although there are many variations, in the most common analysis workflows, proteins extracted from a sample are digested into peptides using a protease, and the resulting peptide mixture is separated by liquid chromatography in a manner that gradually introduces charged peptide ions into a mass spectrometer. As the ions stream in, the instrument measures the *m/z* of these precursor peptide ions, fragments them into many smaller ions, and acquires mass spectra of the ensemble of fragment ions, thereby creating a digital record of the content of each injected sample^2^.

The interpretation of the mass spectra thus produced from each sample requires advanced software to determine putative peptide and protein identifications, confidence metrics for those identifications, and abundance measurements based on the signal intensities^3^. The software available for such processing includes free and open-source packages written by researchers in the community, commercial offerings from the instrument vendors themselves, as well as software tools from independent companies^4^. In typical analysis strategies the spectra are analyzed by matching their peak patterns to a search space of peptide ions that may be present in the sample, either in the form of a database of possibly present protein sequences or a library of previously identified spectra. In both cases, if the exact combination of peptide sequence, amino acid modifications, and charge state is not present in the search space, then the spectrum cannot be correctly identified. Several groups have demonstrated the ability to open the search space to consider unpredicted modifications^5–9^, but these strategies generally lead to an overall decrease in identifications at a given FDR threshold, so are not widely adopted in bottom-up proteomics.

Sequence database searching is still the most commonly used workflow, in which a search engine, such as Comet or X!Tandem, iterates through a list of input spectra, selects from a list of protein sequences a set of peptides that have the same precursor *m/z* within a selected tolerance, and scores each spectrum against a theoretical prediction of the fragments produced from each candidate peptide^10, 11^. The most common format for this protein sequence database is the venerable FASTA format^12^, a simple format that encodes an identifier, a free-text description, and the sequence for each protein. The format is very simple, used by most search engines and downstream processing tools, and is exported by nearly every purveyor of protein sequence lists. In cases where a sequence search engine does not use FASTA, there is a pre-indexing or pre-processing program to transform FASTA files into the needed format.

However, the FASTA format has several widely-recognized shortcomings. First, FASTA files cannot contain metadata about the collection itself: its origin, its production date, key assumptions and parameters used in its production, etc. Second, the description line for each entry is unstructured free text into which different file producers insert entry level metadata in a variety of ways that resists consistent interpretation by reading software packages; even the identifier of a single protein is subject to variations of parsing, making the mapping of proteins across different versions of a FASTA file difficult. Third, there is no mechanism for annotating the locations and nature of known post-translational modifications (PTMs) and sequence variants, which are becoming increasingly important in comprehensive analyses of datasets and to describe actual proteoforms. The UniProtKB/Swiss-Prot .DAT format does allow for encoding of variants and PTMs, but is not standardized or commonly used to inform database searching. A few software packages have custom mechanisms for searching for variants in knowledge bases (e.g. a second, refined search in X!Tandem^13^), but none of the implemented mechanisms are broadly accepted, much less ratified as a standard.

The Human Proteome Organization^14^ (HUPO) Proteomics Standards Initiative^15, 16^ (PSI) has been developing and ratifying community-based standards for over 15 years^17^. The standards developed by the PSI range from formats^18^ for MS input^19^, mass spectrometer output^20^, and output from downstream processing tools^21–25^. As proteogenomics studies become more widespread, interest in PTMs grows, and the available computational capacity expands, the deficiencies in the FASTA format have become an acute problem that would be well remedied with a community-developed enhanced standard from the PSI. All proposed standards are first subjected to the PSI Document Process^26^, a three-level process of review that must be completed before any proposal is declared a ratified standard.

Here we present a new format from the PSI to address the need for an improved FASTA format, the PSI Extended FASTA Format (PEFF). In this article we first present an overview of the format, a brief description of its most salient features, and some example applications. We then describe the available PEFF resources, including the full specification, example files, format validators, software libraries, viewer applications, search engines that implement it, and data providers that already produce it. We finish with a discussion of important applications and considerations for this new format.

## Format Description

The PEFF schema has two main sections as depicted in Figure 1. First is the file header section, which provides metadata about the collection itself, including support for independently describing several source databases that may be merged within one file. This section is absent from FASTA files. In PEFF files, each header line is prefixed with a “#” character (ASCII 35) so that FASTA readers-that are able to ignore comment lines beginning with “#”-can read PEFF files without software changes. In terms of readability, a space following the “#” is preferred, but not mandatory. Second is the individual sequence entries section, which appears in a similar pattern as FASTA files, albeit with more extensive and explicitly constrained annotation.

**Figure 1.**
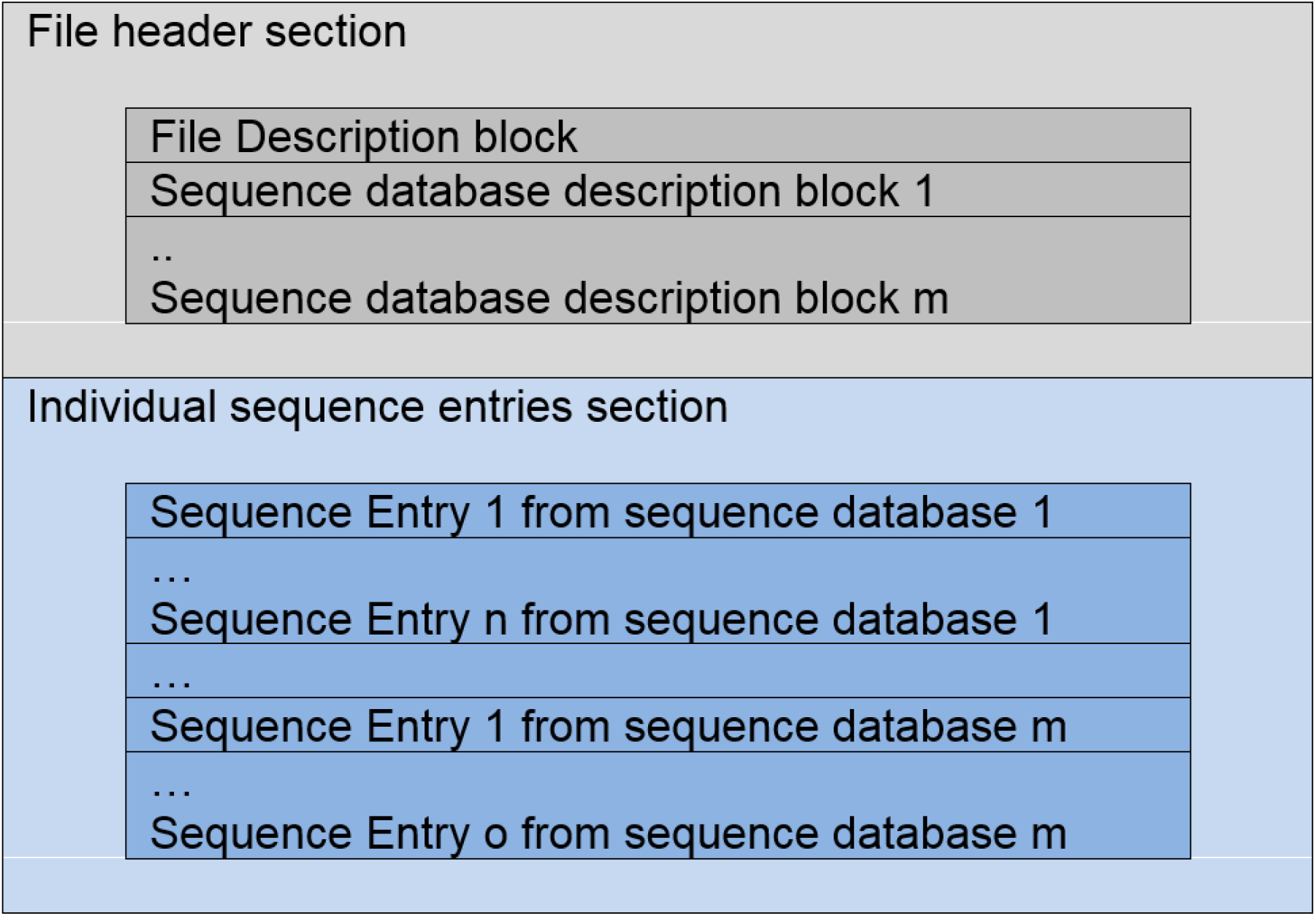
Overview of the PEFF schema. The file header section encodes metadata about the file itself and about the one or more sequence databases contained in the file. The individual sequence entries section encodes each of the individual sequences and the metadata associated with each entry.

A crucial component of the PEFF schema is that a controlled vocabulary is used to specify the permitted keys in the key-value pairs encoded in a PEFF document^27^. This ensures that all values for the same concept are stored under the same key across all PEFF documents, quite unlike FASTA. There is a mechanism for formally defining custom keys to support cases where custom pipelines may wish to implement some non-standard key-value pairs. Custom keys may be tied to concepts in other controlled vocabularies by providing a CURIE (compact URI) to that term. This is generally discouraged for publicly released files, but is available for judicious use. The PEFF controlled vocabulary keywords are stored in a special branch of the main PSI-MS controlled vocabulary^28^ (https://www.ebi.ac.uk/ols/ontologies/ms), which is already widely available and extensively used in extant software and PSI formats. PTMs are encoded in PEFF with entries from the Unimod^29^ or PSI-MOD^30^ controlled vocabularies.

The file header section has three main components. First, the preamble indicates the PEFF format version number. Second, a series of key-value pairs encodes metadata about the origin of the file. Third is a series of one or more key-value pair groups that describes each of the one or more constituent databases in the file. For example, a PEFF file may contain both neXtProt^31^, RefSeq^32^ sequences, and an explicit decoy sequence database in the same file and describe their origins individually.

The individual sequence entries section is essentially the same as in a FASTA file with the two main exceptions that all sequence identifiers must contain a source database prefix as defined in the file header section, and the rest of each description line is constrained to be a series of key-value pairs, where the keys are defined in the controlled vocabulary. This ensures consistent parsing by all readers that properly implement the PEFF specification. Table 1 lists an example (non-exhaustive) set of key-value pairs and their interpretation.

**Table 1.**
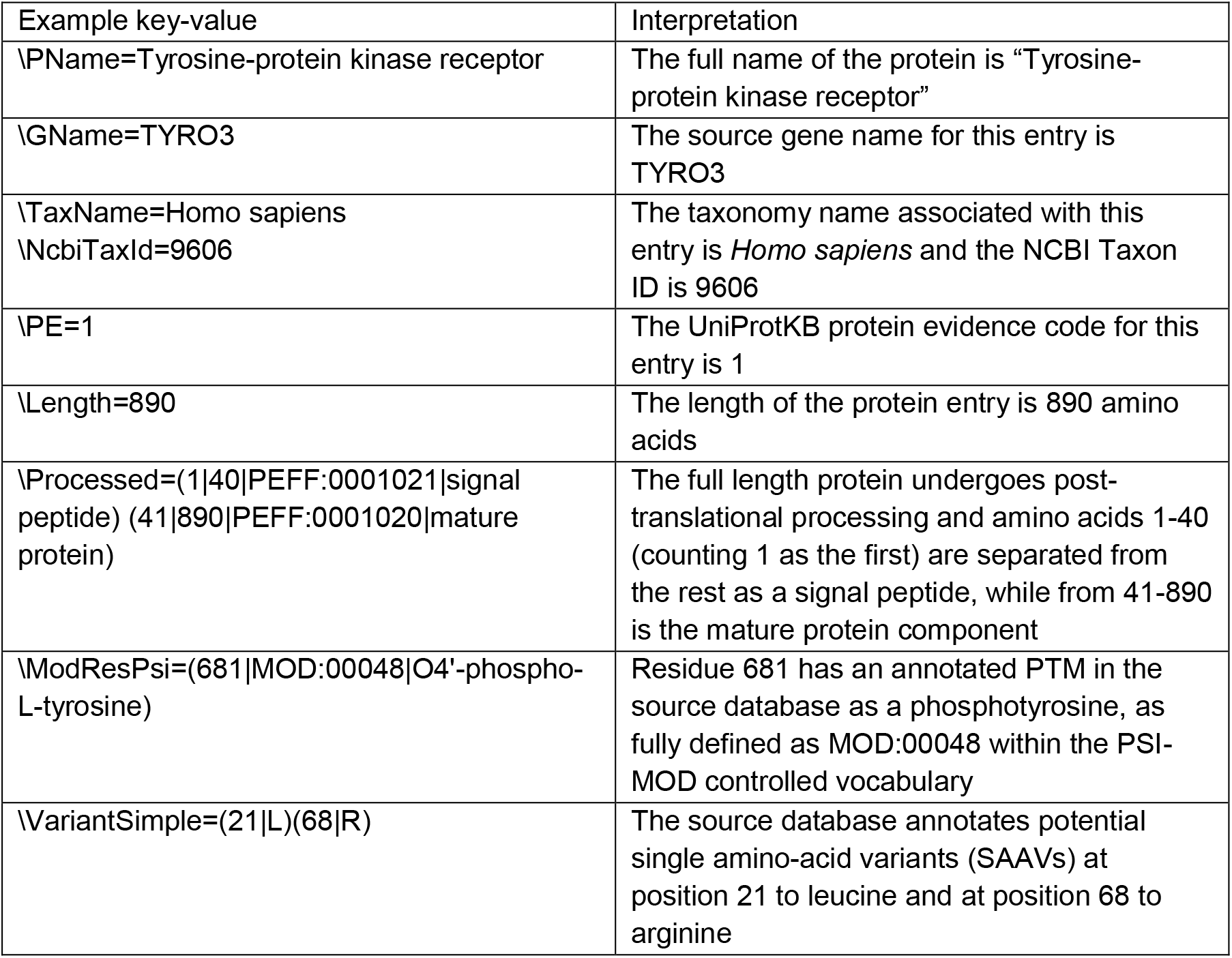
A set of illustrative example key-value pairs that could appear in the description line of a PEFF file. All keys are defined in the PSI-MS controlled vocabulary (https://www.ebi.ac.uk/ols/ontologies/ms).

PEFF is primarily designed to encode a set of reference protein sequences and the associated collection of annotations on each protein, most commonly in the form of potential PTMs and sequence variants. However, any of the constituent databases can be defined in a PEFF header as being a database comprising proteoforms. A proteoform is defined as any one of the multitude of protein forms that can result from a single gene, including sequence variations, PTMs, and processing results^33^. There have been several other efforts to define nomenclatures, ontologies and notations for proteoforms^34, 35^, including the recent ProForma^36^, although the latter focuses more on capturing the results of experimental analysis than being a mechanism for encoding the contents of a protein knowledge base.

There are two methods in which proteoforms can be defined in a PEFF file: the long method, wherein each entry is a different proteoform, and the compact method, wherein each entry defines a basic template and set of interchangeable annotations that may be assembled in different combinations to create multiple proteoforms per entry. In the long method (denoted in each database header via the *isProteoformDb=true* flag), each sequence entry is required to be a single proteoform, where all key-value annotations that describe variation must apply to that sequence. For example, if five PTMs are listed, all are applicable to that specific proteoform entry. Sequence variation-defining key-value pairs are discouraged for proteoforms; however, if supplied, they must be applied. By using this extension, top-down proteomics and other similar applications can create and use a PEFF file of known proteoforms for analysis.

A more compact form is also available via the use of the *hasAnnotationIdentifiers=true* flag in the database header (*isProteoformDb=true* and *hasAnnotationIdentifiers=true* are mutually exclusive in the same database). In this form, as depicted in Figure 2, each sequence entry is a basic template with a set of potential variations, plus a special *\Proteoform* keyword that specifies which of the optional PTMs, sequence variants, disulfide bonds, and processing events should be applied to the template in combination to create individual proteoforms. In this form, the database may be used by ordinary bottom-up applications by ignoring the *\Proteoform* keyword, and also used by top-down applications by automatically expanding the proteoforms based on the listed annotation combinations.

**Figure 2.**
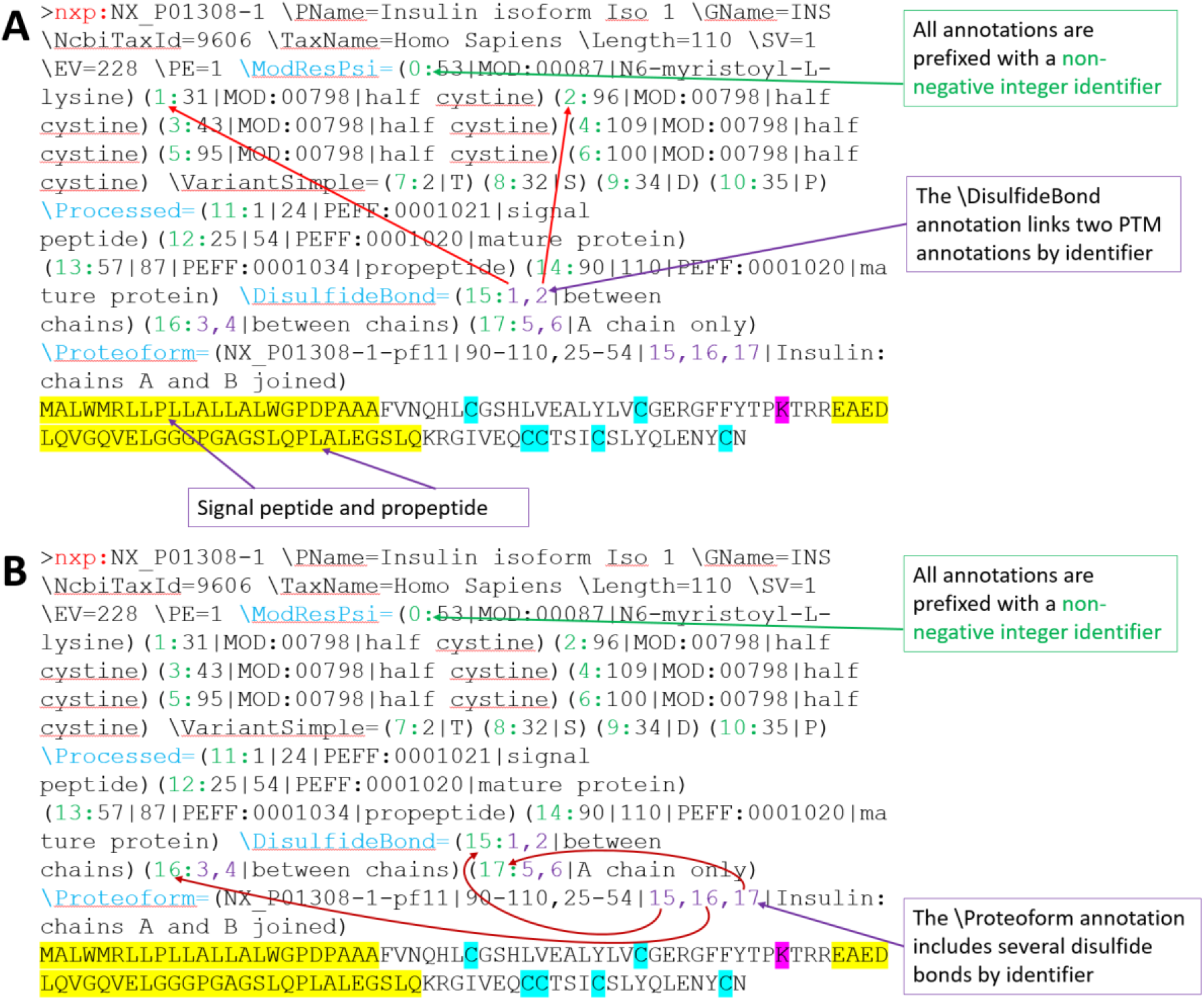
Simplified depiction of how annotation identifiers can be referenced by other annotations to link them, such as for disulfide bonds and for proteoform definitions. Each annotation has a non-negative integer identifier, and other annotations may link to them. This example (somewhat simplified for clarity of presentation) for human insulin encodes: A) PTMs and disulfide bonds that link two PTMs; and B) a final proteoform that include two separate processed chains that are linked together via disulfide bonds.

## Resources and Implementations

There are many components that help define PEFF in addition to this article, which merely provides a brief overview. Further details about PEFF can be obtained at the PSI web page for PEFF (http://www.psidev.info/peff) as well as at the GitHub repository page (https://github.com/HUPO-PSI/PEFF), where version-controlled files are managed.

The primary document is the official PEFF Format Specification (https://github.com/HUPO-PSI/PEFF/tree/master/Specification). This document has been jointly developed by the PEFF designers and subjected to the PSI Document Process in conjunction with many of the additional resources described below, prior to final ratification. The specification document presents all the details needed to implement a PEFF reader or writer successfully.

Accompanying the PEFF Format Specification is a series of example files, including a smallest possible valid PEFF file, a series of increasingly complex but human digestable examples, and a set of invalid files that can be used to test PEFF reading implementations. An important component of the PEFF Format Specification is the PEFF validator, which is able to read a PEFF file and report any warnings or errors on its adherence to the specification. The validator is available as a web application or can be downloaded at http://www.psidev.info/peff.

There is also a Perl library available for download for reading, writing, and modifying PEFF files. The Proteomics::PEFF Perl library comes with a tool that enables easy editing of PEFF files programmatically. For example, it can convert a FASTA file to a PEFF file, and it can add a series of additional PTMs or variants to individual proteins to an existing PEFF file, based on a simple tab-separated list of changes to make. The phpMs^37^ toolkit (http://pgb.liv.ac.uk/phpMs/) also supports the viewing and creation of PEFF files. Pyteomics 4.0^38^, a proteomics software library for the Python language, supports PEFF reading. Implementations in other languages are underway. An up-to-date summary of implementations is available at http://psidev.info/peff.

The neXtProt knowledge base has been exporting PEFF files of its builds since 2015. However, it should be noted that the exports prior to February 2019 did not conform to the final PEFF Format Specification, but rather to earlier draft versions, which are subtly different. This is a natural outcome of the standards development process wherein neXtProt exported their data according to the active draft of the PEFF Format Specification to enable software testing of the format. UniProt^39^ has implemented an export of its variation data using PEFF via the Proteins API^40^ (https://www.ebi.ac.uk/proteins/api/doc/).

The ultimate utility of PEFF will be in its implementation in proteomics search engines and downstream analysis and visualization software. As of this writing, the Comet search engine^41^ has been adapted to read PEFF files (in addition to FASTA files) and process input MS data using the encoded variants and PTMs. The Trans-Proteomic Pipeline^42–44^ (TPP) will soon implement PEFF in its downstream validation and visualization of data searched with Comet using PEFF input. The ProteoMapper^45^ tool (http://www.peptideatlas.org/map/) can search a PEFF file for a set of input peptide sequences, taking into account the protein variations encoded in PEFF. Submission of datasets to ProteomeXchange^46, 47^ supports the inclusion of the reference database used. Currently this usually means FASTA files; going forward, PEFF files should be similarly submitted or cited when they are used as a reference. A complete summary of supporting software and resources is available and will be maintained as tables of producers and consumers of PEFF at http://www.psidev.info/peff.

## Discussion and Conclusions

The choice to expand on the basic structure of the FASTA format has not been made without dissenting opinions during the design of PEFF. Porting an existing FASTA parser to a PEFF parser will be quite easy for the most basic features. However, as the more advanced features of PEFF are parsed, the job of parsing a complex free-text format becomes considerably more difficult. Alternative encoding strategies such as a single XML (Extensible Markup Language) file and a side-car annotations file that is separate from a FASTA file were seriously considered. Parsing of complex sequence annotations from a PEFF-like XML format in general would be easier via the use of existing XML-parsing frameworks, but this requires completely new parsers and additional software dependencies. The PSI philosophy over the years has generally been to avoid side-car implementations since these types of files have a tendency to become separated from their siblings, thus causing information loss. In the end, the predominating opinion that PEFF should retain the FASTA format’s basic structure and thereby should enable a modest upgrade path for existing FASTA parsers rather than require completely new parsers prevailed.

Standard file formats are only as effective at the software that implements them. However, this precept can often be a chicken-and-egg problem in that it is often difficult to finalize a standard until it has been well tested by several implementations, and yet it is difficult to convince software developers to implement a format that has not yet been finalized. PEFF has finally achieved critical mass with one major search engine implementation (Comet) several major exporters (neXtProt and UniProt) supporting PEFF, and emerging research citing the use of PEFF in the workflow^48^. As a key point, several software libraries now support PEFF. Additionally, the Protein Prospector^6^ search engine is currently in the process of implementing PEFF support (after previously supporting similar functionality with *ad hoc* formats). Therefore, we expect the number of implementing resources to expand rapidly once PEFF has been ratified by the PSI.

One of the driving applications for PEFF is proteogenomics^49^, in which the variations unique to each sample from each distinct individual are important to the data analysis. In such scenarios, genomic sequencing, RNA-seq, or ribosome profiling (e.g., using PROTEOFORMER^50, 51^; https://github.com/Biobix/proteoformer) will determine the variations unique to the sample, and that information will be used to create a custom sequence database specifically for that sample. PEFF provides an ideal format for this workflow. PEFF provides support for analysis workflows where nucleotide sequences are used as the primary sequence information. Each database within a PEFF file can be defined as an amino acid database or a nucleotide database. Molecule type can be mixed within a file, but not within one database. It is similarly intended that PEFF will enable top-down analysis workflows, as we better understand the full complement of proteoforms detectable in biological samples. In this context, the previously mentioned notation ProForma^36^ has been recently developed by the Top Down Proteomics Consortium. Proforma uses a different style of notation that embeds the annotations into the sequence. We have not incorporated this format into the PEFF sequences component, since the proteoforms can equally be described in the PEFF format, and it is preferable not to offer several ways to encode the same information, since this increases the complexity for parsers.

The PSI is an open consortium of interested parties, and we encourage participation and critical feedback, suggestions and contributions to PEFF and other PSI formats *via* participation at PSI annual workshops, conference calls, the GitHub collaboration platform, and PSI mailing lists (see http://www.psidev.info/).

## Acknowledgements

We thank the many contributors to early versions of this format, including David Creasy, Matt Chambers, Phil Andrews, members of the UniProt Consortium, particularly Nicole Redaschi, Maria Jesus Martín, Claire O’Donovan, Peter McGarvey, and Amos Bairoch for mapping the proposal with UniProtKB concepts. We would like to thank Frédéric Nikitin, Pierre-André Michel, Monique Zahn and the whole neXtProt team at SIB for the PEFF implementation in neXtProt. We also want to thank Edward Turner, Leonardo Gonzales, Guoying Qi, Andrew Nightingale and Jie Luo from the UniProt development team at EMBL-EBI for the PEFF implementation in the Proteins API. EWD is supported in part by the National Institutes of Health (NIH) grants R24GM127667, R01GM087221, U19AG023122, and U54EB020406. LFHS is supported by the European Research Council and the Research Council of Norway. HB is supported by the Research Council of Norway and the Bergen Research Foundation. LL is supported by the SIB Swiss Institute of Bioinformatics. EA is supported by the NIH under the UniProt grant U24HG007822 and by the European Molecular Biology Laboratory (EMBL). GM is funded by the BMBF grant de.NBI - German Network for Bioinformatics Infrastructure (FKZ 031 A 534A). The funding of ME is related to PURE and VALIBIO, projects of Northrhine-Westphalia. JAV wants to acknowledge funding from EMBL, the Wellcome Trust (grants WT101477MA and 208391/Z/17/Z) and NIH (grant R24GM127667). RJC and PRB are supported by the Adelson Medical Research Foundation.

## Author Information

The authors declare no competing financial interest.

## Supporting Information

None

